# NMDtxDB: Data-driven identification and annotation of human NMD target transcripts

**DOI:** 10.1101/2024.01.31.578146

**Authors:** Thiago Britto-Borges, Niels Gehring, Volker Boehm, Christoph Dieterich

## Abstract

The nonsense-mediated RNA decay (NMD) pathway is a crucial mechanism of mRNA quality control. Current annotations of NMD substrate RNAs are rarely data-driven, but use general established rules. We introduce a dataset with 4 cell lines and combinations for SMG5, SMG6 and SMG7 knockdowns or SMG7 knockout. Based on this dataset, we implemented a workflow that combines Nanopore and Illumina sequencing to assemble a transcriptome, which is enriched for NMD target transcripts. Moreover, we use coding sequence information from Ensembl, Gencode consensus RiboSeq ORFs and OpenProt to enhance the CDS annotation of novel transcript isoforms. 302,889 transcripts were obtained from the transcriptome assembly process, out of which, 48,213 contain a premature stop codon and 6,433 are significantly up regulated in three or more comparisons of NMD active vs deficient cell lines.

We present an in-depth view on these results through the NMDtxDB database, which is available at https://shiny.dieterichlab.org/app/NMDtxDB, and supports the study of NMD-sensitive transcripts. We open sourced our implementation of the respective web-application and analysis workflow at https://github.com/dieterich-lab/NMDtxDB and https://github.com/dieterich-lab/nmd-wf.

## Introduction

Eukaryotic gene expression is a regulated process that culminates in the production of a messenger RNA (mRNA) that serves as a template for protein synthesis. Pre-mRNA undergoes co-transcriptional processes like capping, splicing, and polyadenylation before becoming mature mRNA (1). In the cytoplasm, mature mRNAs are translated into proteins by ribosomes. In this context, quality control mechanisms ensure mRNA production fidelity, and the nonsense-mediated mRNA decay (NMD) is central to this process. This pathway identifies and degrades mRNAs with premature termination codons (PTC), hence preventing translation of truncated protein products. Dysregulation of the pathway has been extensively implicated in deleterious cellular outcomes and, consequently, human disease (2,3). PTC can arise from various causes, including DNA mutations or alternative splicing events (4,5). Despite the relevance of this pathway to RNA biology, NMD transcripts representation in genome annotation projects, such as the Ensembl database, is incomplete.

There are two mechanistic models that describe the activation of NMD, with the exon-junction complex (EJC)-dependent model being the most extensively studied mechanism. The EJC is a protein complex deposited around 22 nucleotides (nt) upstream of exon-exon junctions after splicing. During translation, all EJCs are normally displaced from the transcript by the ribosome, since stop codons are usually located in the last exon. However, when ribosomes terminate translation at a PTC, which is defined as being located more than 50 nucleotides upstream of the last exon-exon junction, any downstream EJC(s) remain bound on the transcript. Consequently, the presence of residual EJC components downstream of terminating ribosomes is a strong determinant for activating NMD. This activation involves the SMG1-dependent phosphorylation of UPF1, which is the starting point for the decay process. Hyperphosphorylated UPF1 engages with both the SMG6 and SMG5/SMG7 complexes. SMG5/SMG7 are needed to access the endonuclease activity of SMG6, which cleaves the mRNA near the PTC site, allowing for rapid degradation of the decay intermediates by exonucleolytic factors. Therefore, NMD activation results in rapid mRNA decay and depletion of potentially toxic RNA entities.

An alternative pathway for activating NMD, referred to as the “faux 3′ UTR” model (6) requires mRNAs with a longer (over 400 nt) 3’ untranslated region (3’ UTR). This extended 3’ UTR allows for higher accessibility to UPF1 association. However, the mechanistic properties of this mechanism are still actively debated in the field. Not all mRNAs possessing long 3’ UTR are inevitably subjected to decay via NMD, and binding of specific RBPs, such as hnRNP L, PTBP1 or PABPC1, can counteract the process (7).

The rules driving the EJC-dependent activation of NMD have been determined with mechanistic detail using reporter gene constructs. As stated above, the primary rule states that an EJC situated more than 30-35 nucleotides downstream of a translation termination codon triggers NMD (8). This basic 50nt rule, representing the distance of the last exon-exon junction to the PTC, is currently applied in databases such as Ensembl to annotate transcripts and genomic variants that trigger or escape the pathway.

Genome annotation projects such as Ensembl are the primary source for the annotation of NMD targets. However, these sources are insufficient and do not keep pace with the recent surge in the discovery of novel transcript isoforms identified by RNA long-read technologies. While databases housing information on nonsense mutations are widely available, databases for splicing events, which may lead to NMD activation are rare. Furthermore, even though the number of datasets used to study NMD depleted conditions is increasing, obtaining specific isoforms that are targeted by NMD is a challenging task.

Here, we present our detailed transcriptome analysis using short and long-read sequencing of 4 human cell lines across control and conditions, which deplete three NMD key factors: SMG5, SMG6 or SMG7. In addition, we introduce a computational workflow to predict NMD-sensitive transcripts from these libraries, and annotate their PTC status. Finally, we integrate these results in a database, NMDtxDB, which is equipped with an intuitive web interface that allows researchers to interpret the NMD status of a given transcript in context to the expression and transcript structure features.

## Results

### Datasets and data resources

NMDtxDB comprises data from 54 RNA-libraries. The libraries were derived from 4 cell lines, and were sequenced by Illumina short-read sequencing in biological triplicates, comprising controls and samples depleted for three key NMD factors. A subset of 8 libraries, 4 biological replicates for treatment and control conditions for the HEK293 SMG7 KO SMG6 KD comparison, were additionally sequenced with Nanopore Direct RNA long read sequencing (DRS). The dataset is described in detail in Table 1.

**Table 1.** Description of the RNA-seq datasets used in NMDtxDB. The comparison with an asterisk (HEK_SMG7KO_SMG6KD_Z245*) was additionally sequenced with Nanopore Direct RNA Sequencing. KD were compared against a Luciferase (Luc) siRNA for the same cell type, genotype (WT or SMG7 KD), and clone. The study has a treatment (NMD factor KD) vs control (Luc KD) design, and samples were sequenced in three (Illumina) or four (Nanopore DRS) biological replicates.

| Comparison | Cell line | siRNA KD | CRISPR KO |
| --- | --- | --- | --- |
| HEK_NoKO_SMG5KD_Z023 | HEK 293 | SMG5 |  |
| HEK_SMG7KO_SMG5KD_Z245 | HEK 293 | SMG5 | SMG7 |
| HEK_SMG7KO_SMG6KD_Z245* | HEK 293 | SMG6 | SMG7 |
| HEK_SMG7KO_SMG5KD_Z319 | HEK 293 | SMG5 | SMG7 |
| HEK_SMG7KO_SMG6KD_Z319 | HEK 293 | SMG6 | SMG7 |
| HEK_NoKO_SMG6+SMG7KD_Z023 | HEK 293 | SMG6+SMG7 |  |
| HeLa_NoKO_SMG6+SMG7KD_Z021 | HeLa | SMG6+SMG7 |  |
| U2OS_NoKO_SMG6+SMG7KD | U2OS | SMG6+SMG7 |  |
| MCF7_NoKO_SMG6+SMG7KD | MCF7 | SMG6+SMG7 |  |

### De novo transcriptome assembly

We developed a reference-guided transcriptome assembly workflow using StringTie (12) to detect transcript isoforms present in NMD-depleted conditions. The reconstruction of RNA isoforms based on RNA-seq reads, and reference transcript assembly poses several challenges that affect the quality of the resulting transcriptomes. The first challenge is the relatively short length of read fragments obtained with Illumina sequencing. A second challenge is establishing parameters that eliminate potentially cryptic transcripts, transcripts that are usually associated with low abundance. In the following, we describe how we approach these challenges.

Nanopore Direct RNA-seq data improves the guided assembly approach in two key aspects. It provides longer sequencing reads that allow for the reconstruction of multiple exon-exon chains. Second, it improves the confidence in calling lowly abundant transcripts because it produces a second and complementary view of the transcriptome.

Additionally, we have optimized transcriptome assembly parameters using the CHESS database (13). CHESS was compiled from thousands of RNA-seq samples from the GTEx consortium (14). We provide respective details in the Methods section.

As a result of the transcriptome assembly workflow across 4 cell lines and multiple conditions, we compiled a transcriptome with 302,889 transcripts in 58,006 genes. Of note, 71,114 (23.5%) transcripts have support from Nanopore DRS of HEK293 cells.

In NMDtxDB, we categorized transcripts into four classes based on GffCompare class codes using Ensembl genome GRCh38.p13 v102 as reference (15). Transcripts that have an intron chain match (class code =, *same_intron_chain*) are considered as reference transcripts. All other transcripts are considered novel. Splicing variants share at least one exon-exon junction with reference transcripts, indicating variations in isoform structure (class codes c, j or k). Transcripts with an intron retention event in comparison to a reference transcript were annotated as IR (code n). Transcripts matching to any other class code were classified as other, and are potentially cryptic isoforms. This classification system provides insight into the transcriptome’s complexity of cell lines depleted for NMD factors. The distribution of transcript class codes per long-read (LR) support is shown in Figure 1.

**Figure 1.**
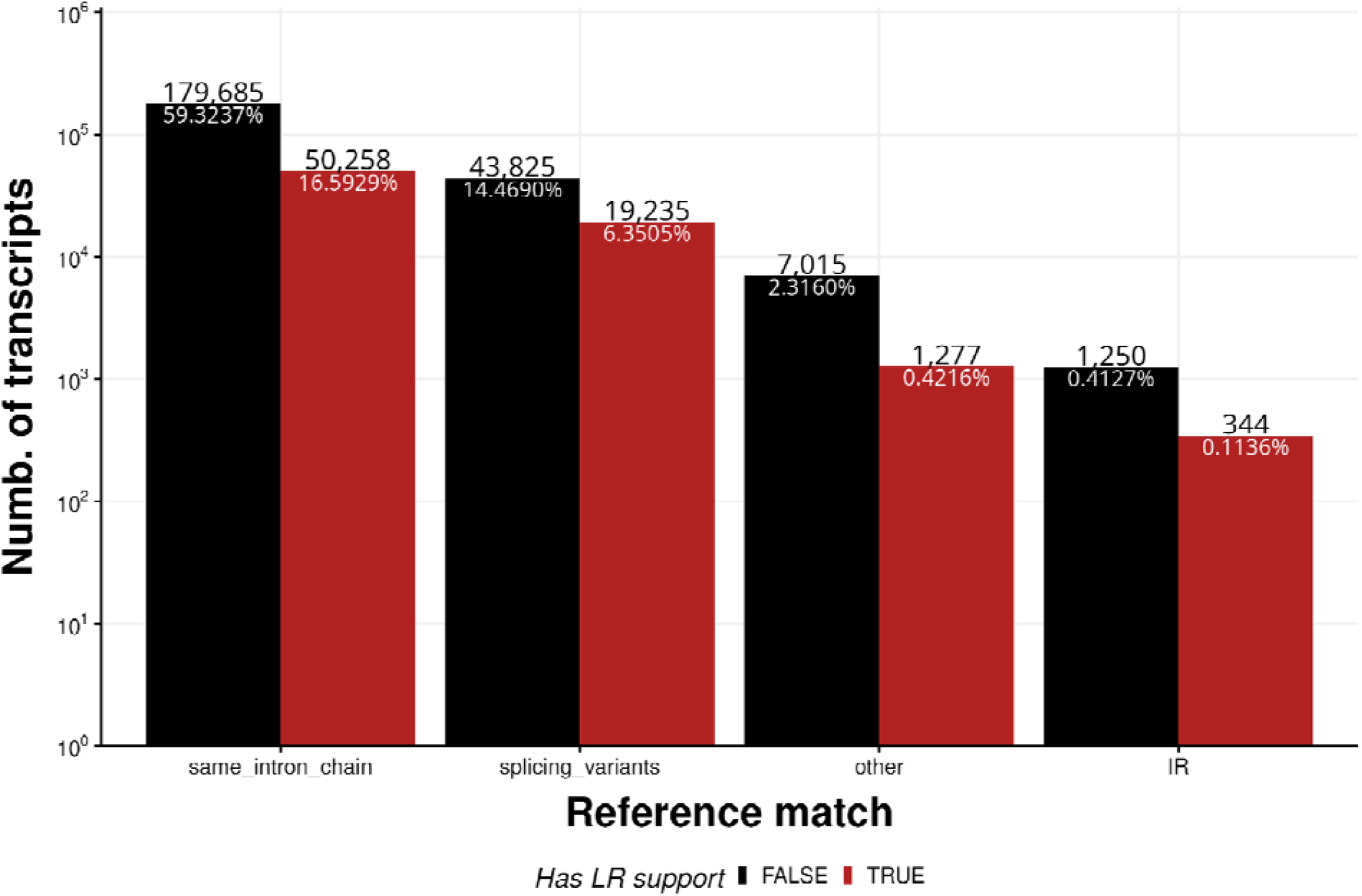
Distribution of transcript per by reference match and long read support. The majority of transcripts (75.9%) match the reference annotation, followed by transcripts that are splicing variants of known transcripts, other transcripts that don’t share features with known transcripts and transcripts that present intron retention.

### Identification of coding sequences and PTC status

We annotate stop codon positions as prerequisite for determining its PTC status. Our methodology involves the integration of three sources of coding sequence information (CDS), which are listed in Table 2. The corresponding workflow is presented in the Methods section. Overall, 102,796 (33.9%) transcripts belong to the *same_intron_chain* class and match to an annotated CDS. We annotated other transcripts classes by start codon projection on transcripts from at least one of the sources in Table 2. Overall, this strategy resulted in CDS annotations for 139,438 (46.0%) transcripts.

**Table 2.** CDS databases used in NMDtxDB. Data from three databases end into four for CDS sources in NMDtxDB. This process is detailed in the methods section, refer to “Coding sequence curation”.

| Database name | Name of NMDtxDB source | Description | Citation |
| --- | --- | --- | --- |
| Ensembl | Canonical, ensembl | Uses a comprehensive process to annotate CDSs. The process has both manual and automated steps. It is the preferred choice for transcriptome annotation. | (15) |
| Encode Ribo-seq ORFs | riboseq | Identifies and catalogs non-canonical, unannotated translated open reading frames in 7 Ribo-Seq datasets. | (17) |
| OpenProt | openprot | Identifies known proteins, novel isoforms, and alternative proteins from publicly available proteomics datasets, and backs up its predictions with conservation, translation, and expression evidence. | (16) |

We then applied the 50 nt rule to predict PTC status after matching transcripts to putative CDS (8). Based on this approach, we detected PTCs in 48,213 transcripts. The breakdown of the PTC per coding sequence is detailed in Figure 2. In addition, we observe a total of 9,419 of PTC transcripts with LR support

**Figure 2.**
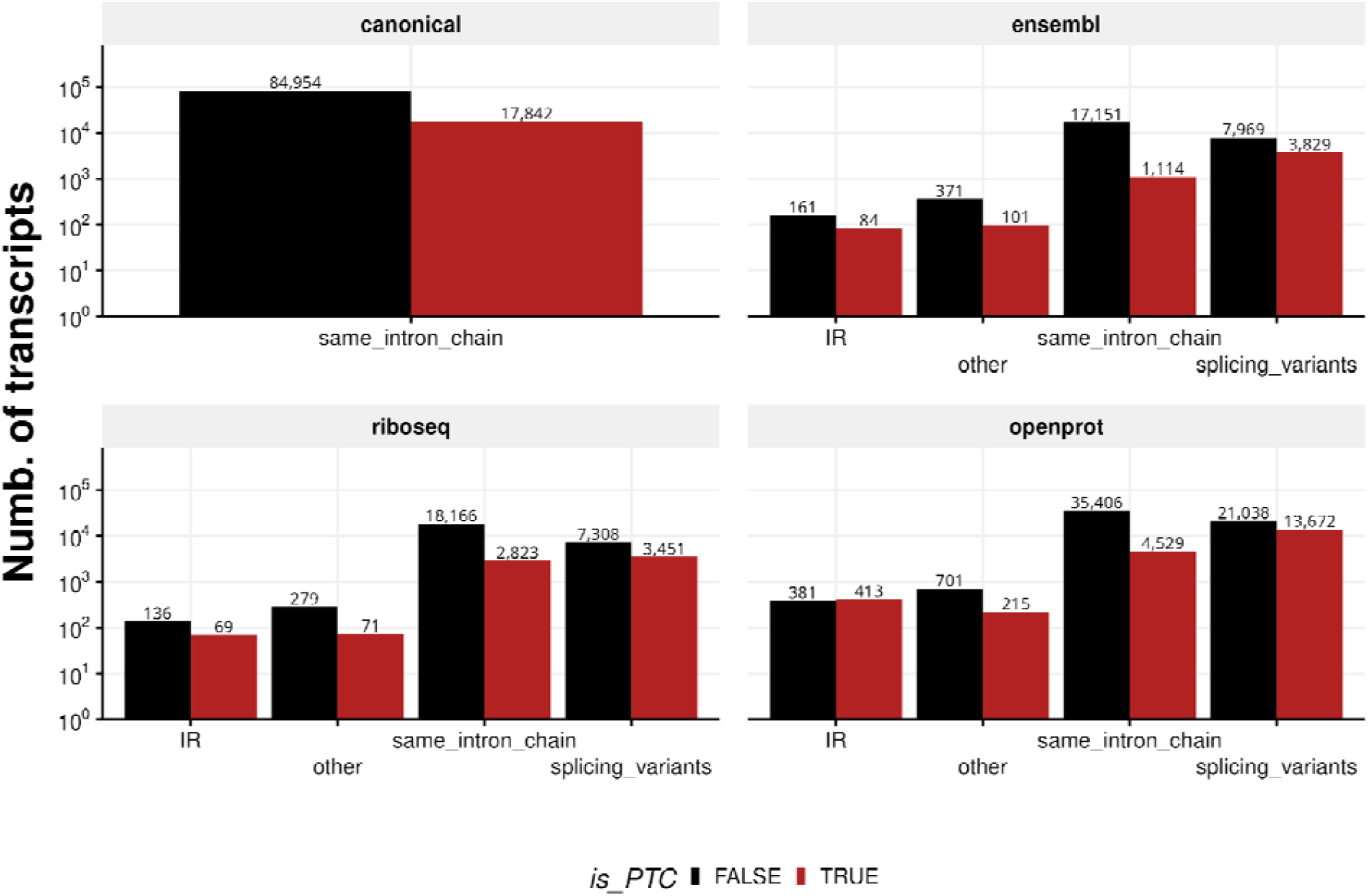
Overview of the transcript-to-CDS matches in NMDtxDB. If transcripts in the transcriptome match to mRNA in the Ensembl annotation, we name these matches as canonical. For novel transcripts or not annotated as mRNA, we use the start codon information from the sources to find a CDS. These may provide multiple CDS per transcript, however, if the CDS is identical, we keep a single hit. The is_PTC flag is TRUE if the distance from the stop codon and the last EJC exceeds 50 nt. Of note, the transcripts in the same_intron_chain class has proportionally less PTC calls for all sources. In contrast, the splicing_variants class has a similar number of transcripts with and without PTC for the riboseq and openprot sources.

### Differential gene and transcript expression analysis

Besides PTC status, another feature that distinguishes NMD targets is differential expression under NMD factor depletion. NMDtxDB provides this information both at the gene and transcript level. For each contrast and gene or transcript, the expression of that entry is compared to the global expression bar plot and colored accordingly. Based on the transcript filtering criteria described in the Methods section, 16,812 genes were tested for differential gene expression (DGE). Figure 3 shows a meta analysis of the DGE comprising the 9 comparisons based on MetaVolcanoR (<u>10.18129/B9.bioc.MetaVolcanoR</u>). Colored dots represent the top 1% genes, ranked by the TopConfect method (18), highlighting that the majority (80.0%) of regulated genes are up-regulated. Out of the 2,177 genes that were called significantly DGE (adjusted p-value < 0.05 and |L2FC| > 1), 1,266 have a sign consistency higher than |5|, meaning the effect size direction was consistent on 6 or more comparisons. Up-regulation of NMD target transcripts in NMD-negative conditions largely contributes to the observed asymmetric gene regulation shown in Figure 3.

**Figure 3.**
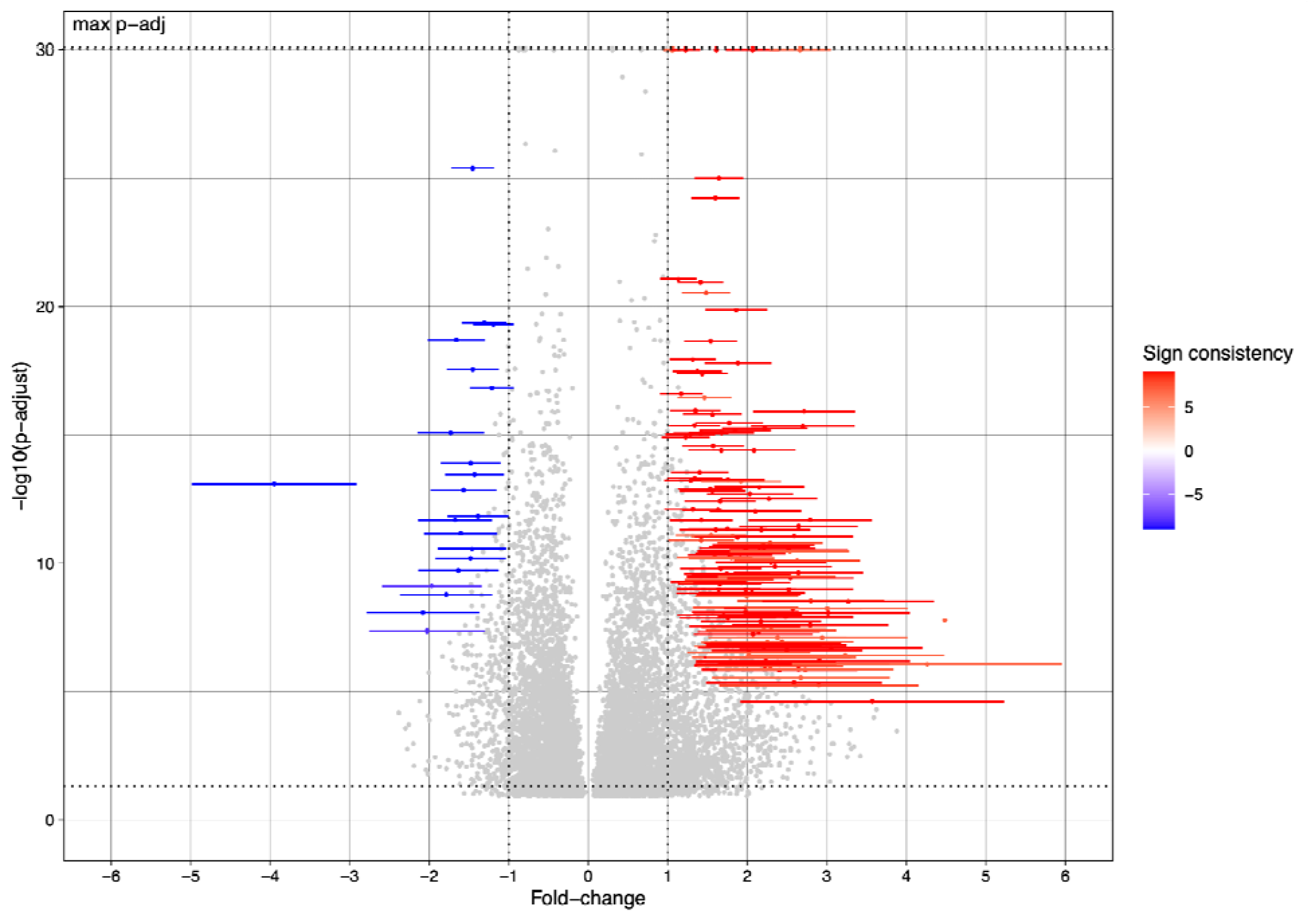
Meta analysis of the differential gene expression. The analysis of gene expression was conducted using the MetaVolcano package, wherein the Fisher’s method and Random Effect Model were employed to summarize each gene’s adjusted p-value and L2FC, respectively, for the 9 contrasts. The top 1% genes, ranked by the TopConfect method, were colored, and the color map represents the direction of change and degree of consistency obtained for the effect size of each gene. This approach summarizes all relevant comparison (NMD vs control) in the dataset. Genes with -log10 adjusted p-value > 30 were set to that threshold to improve visualization.

Table 3 details the number of genes and transcripts called significant for every single comparison. A total of 82,799 transcripts were tested for differential transcript expression (DTE) based on filtering aiming to increase FDR (19).

**Table 3.** Number of DGE and DTE calls in NMDtxDB. The table lists the number of genes and transcripts with DGE and DTE based on cutoff for the adjusted p-value < 0.05 and |L2FC| > 1.

| Contrasts | DGE | DTE |
| --- | --- | --- |
| HEK_NoKO_SMG5-KD_Z023 | 72 | 1,033 |
| HEK_NoKO_SMG6+SMG7-KD_Z023 | 4,856 | 10,105 |
| HEK_SMG7KO_SMG5-KD_Z245 | 3,433 | 13,576 |
| HEK_SMG7KO_SMG5-KD_Z319 | 3,105 | 12,318 |
| HEK_SMG7KO_SMG6-KD_Z245 | 5,215 | 9,870 |
| HEK_SMG7KO_SMG6-KD_Z319 | 4,738 | 9,142 |
| HeLa_NoKO_SMG6+SMG7-KD_Z021 | 4,888 | 12,997 |
| MCF7_NoKO_SMG6+SMG7-KD | 5,236 | 8,506 |
| U2OS_NoKO_SMG6+SMG7-KD | 9,354 | 9,897 |

**Table 4.**
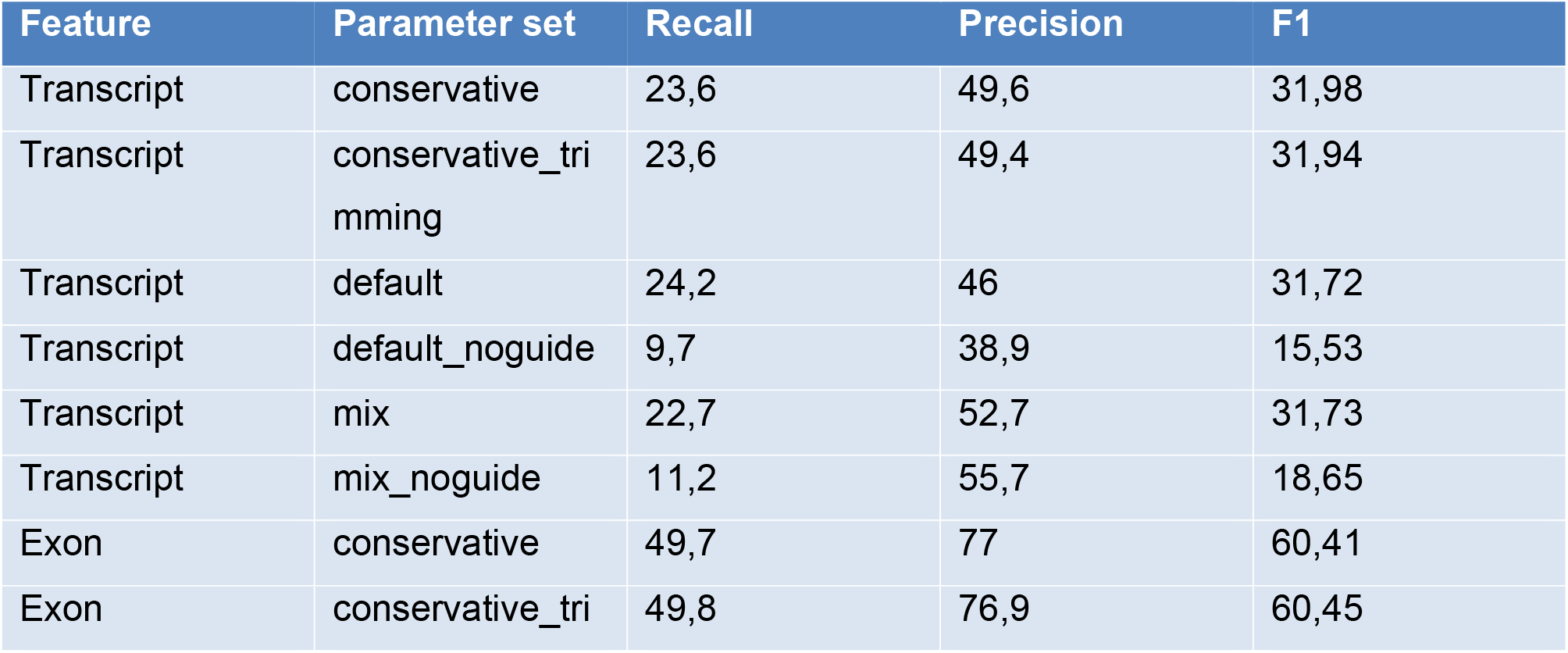
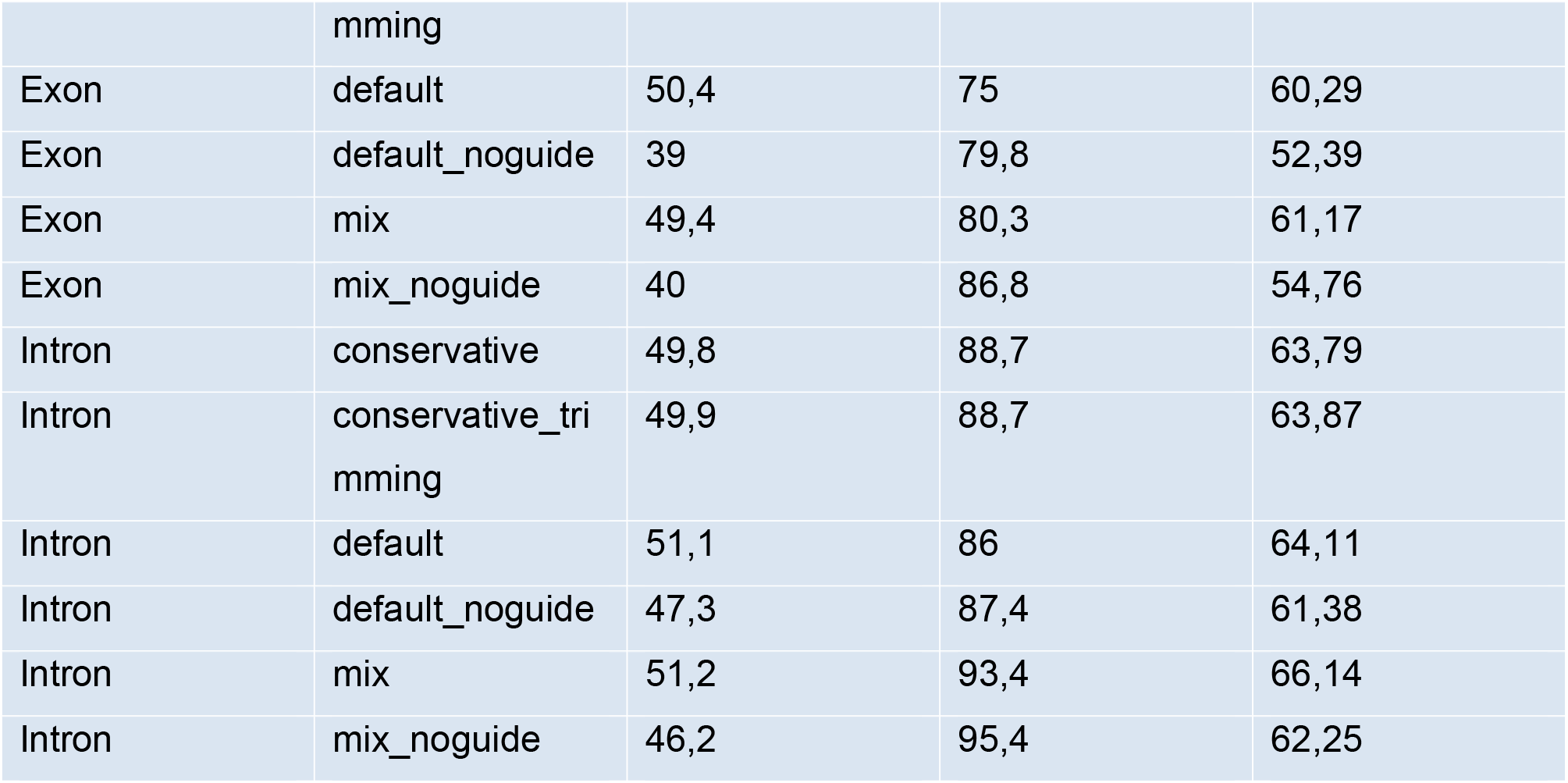
Benchmark of sample 33G-30 versus CHESS. Parameter sets: default: StringTie with default parameters. noguide: StringTie run without reference. Conservative: uses the --conservative flag (same as -t -c 1.5 -f 0.05). trimming: enables trimming (-t flag). mix: uses mix for combining Illumina and Nanopore reads. Bold entries represent the top metric for each feature group.

Next, we integrated the DTE effect sizes with the predicted PTC status to further explore the significance of the 50nt rule based on our annotation pipeline. We considered all information from the aforementioned CDS sources and call a transcript PTC-containing if at least one source indicated that a PTC occurred. Figure 4 illustrates the differential expression patterns of transcripts as stratified by PTC status. Consistent with our expectations, cell lines subjected to NMD factor depletion exhibit a marked up regulation of transcripts that are annotated with a PTC. This is evidenced by a right shift in the median effect size of PTC-harboring transcripts relative to their PTC-lacking counterparts. Notably, the plot also conveys the impact of NMD depletion across the various conditions examined. In particular, the HEK cell lines with SMG5 knockdown (clone Z023) display the smallest effect, as indicated by the proximity of the median expressions for the two groups of transcripts

**Figure 4.**
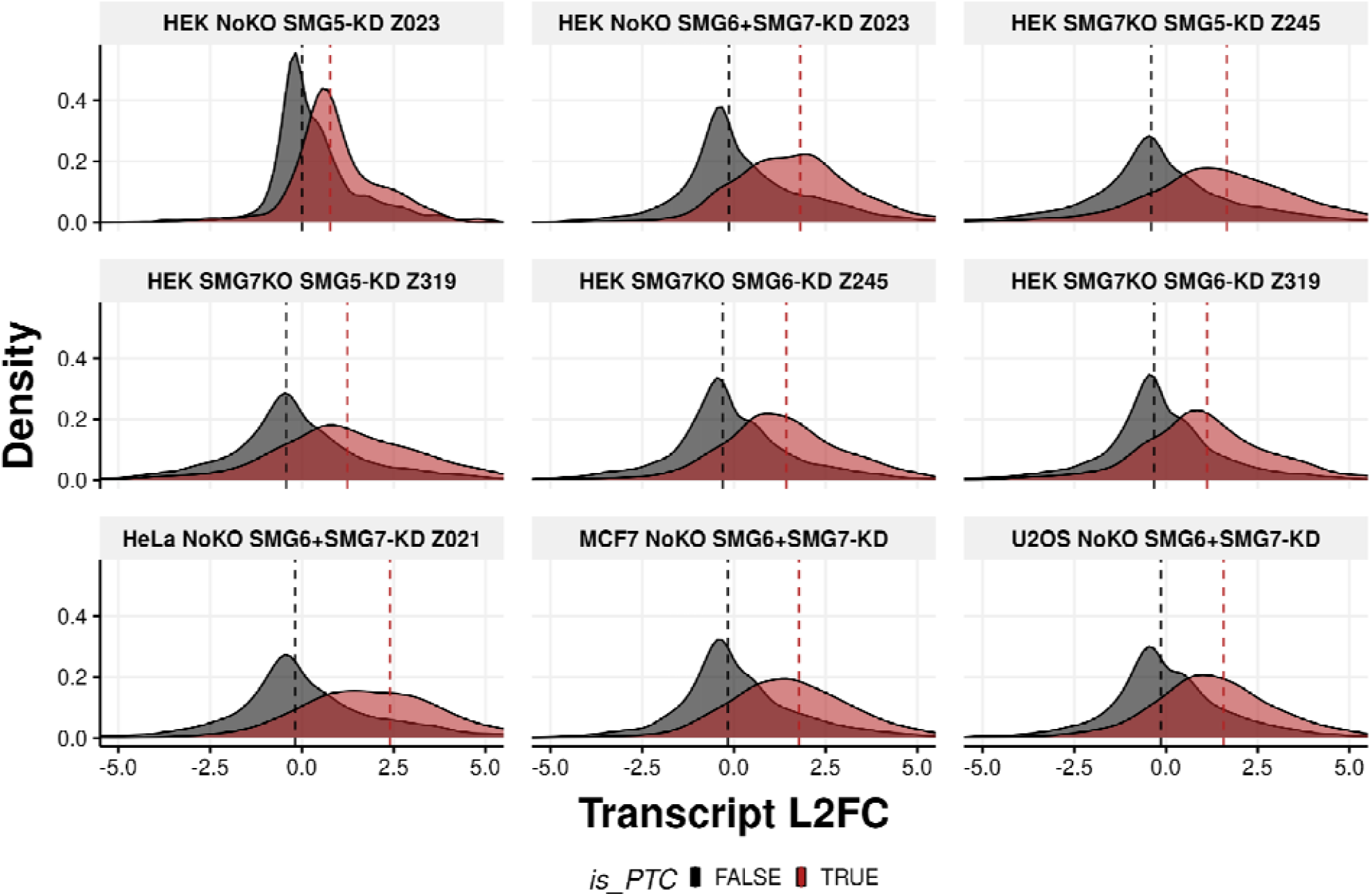
Distribution of differential transcript expression effect size by PTC status. Each facet shows for each comparison the distribution of L2FC (x-axis) and the density (y-axis) for two groups of transcripts (PTC-harboring or not). Only transcripts with a CDS match and adjusted p-value < 0.05 for the DTE were shown. Transcripts were colored by any evidence of PTC, based on any PTC annotation for multiple CDS. Dashed vertical lines represent the median L2FC grouped by PTC status.

### NMDtxDB web-interface

All the aforementioned results are accessible via the NMDtxDB web-interface, which is simple to use, designed for a broad audience, and enables fast information retrieval. The interface is organized into two sections, a sidebar that harbors the selection logic and a main panel that offers either a gene- or transcript-centric view. Moreover, links to Ensembl and Uniprot databases, as well as the UCSC genome browser, are available. Figure 5 shows transcript-specific information on *SRSF2* as an example.

**Figure 5.**
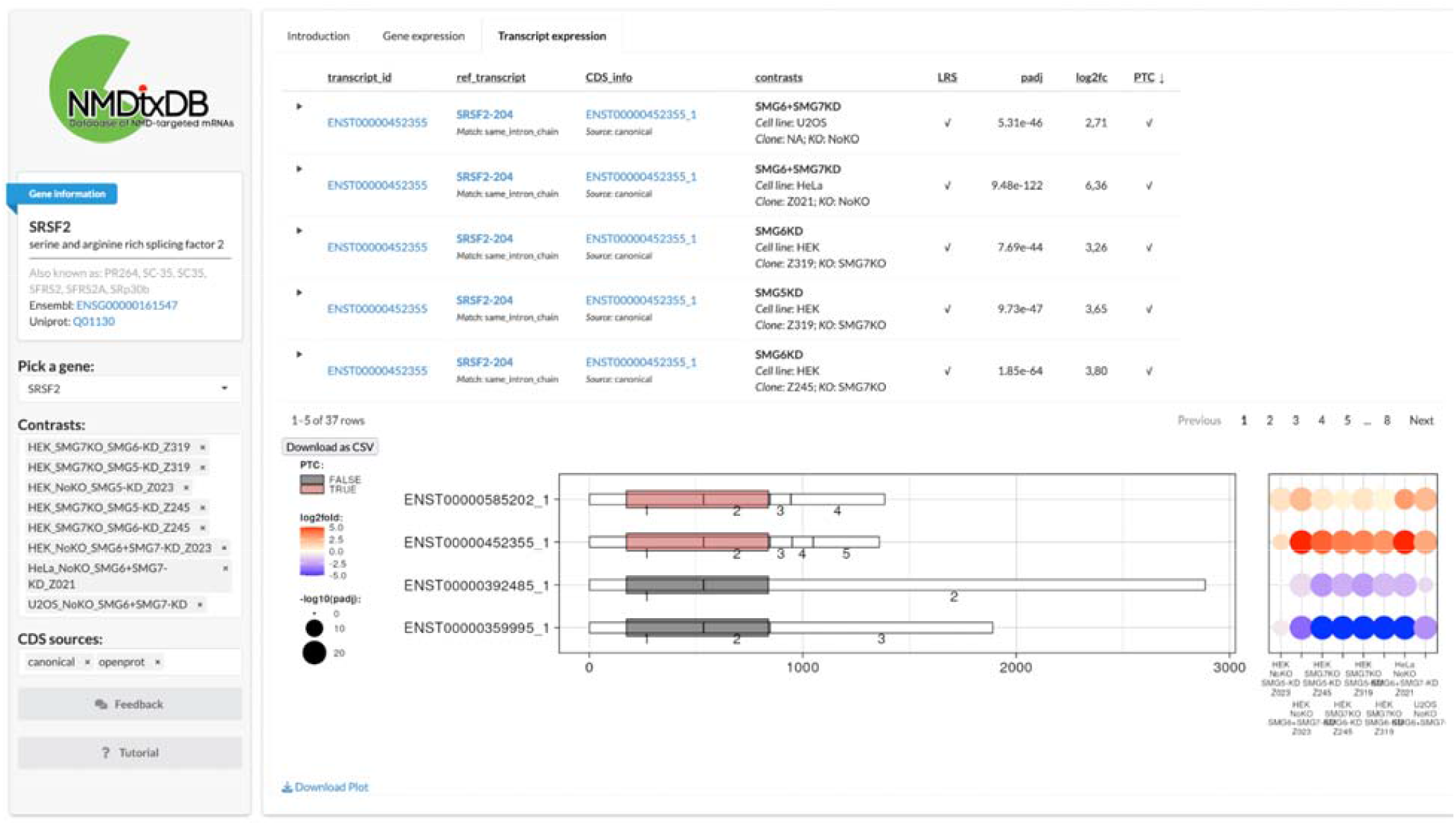
NMDtxDB transcript expression view. On NMDtxDB each view is unified by gene. The sidebar comprises the Gene Information card, followed by the user input for gene, contrast and CDS source. The main panel contains two elements. On the top, a dynamic table listing each transcript-to-CDS match, one entry for each selected contrast, if the transcript was selected for the DTE test. On the bottom, a plot showing the structure for each transcript-to-CDS pair, and a dot plot showing L2FC and p-adjust for each selected contrasts. Each element of th user interface is detailed by the tutorial, and column headers show description by hovering the mouse pointer over them. In this example, we show *SRSF2*, a well described NMD target. Briefly, two transcripts SRSF2-20 (ENST00000452355) and SRSF2-208 (ENST00000585202) are up regulated in 6 comparisons. The transcript structure without introns shows the relative CDS position, which indicates which exons and exon-exon junctions are downstream of the stop codon.

We also provide gene annotation information in NMDtxDB as genome browser tracks. Links within the application point to the UCSC Genome Browser website and automatically load the NMDtxDB TrackHub. This hub consists of the CDS annotation and PTC status for transcripts-to-CDS matches. CDS sources were grouped together for an easier comparison. In addition, coverage tracks are available for a subset of the samples in the dataset. Figure 6 exemplifies the TrackHub view for the *SRSF2* gene.

**Figure 6.**
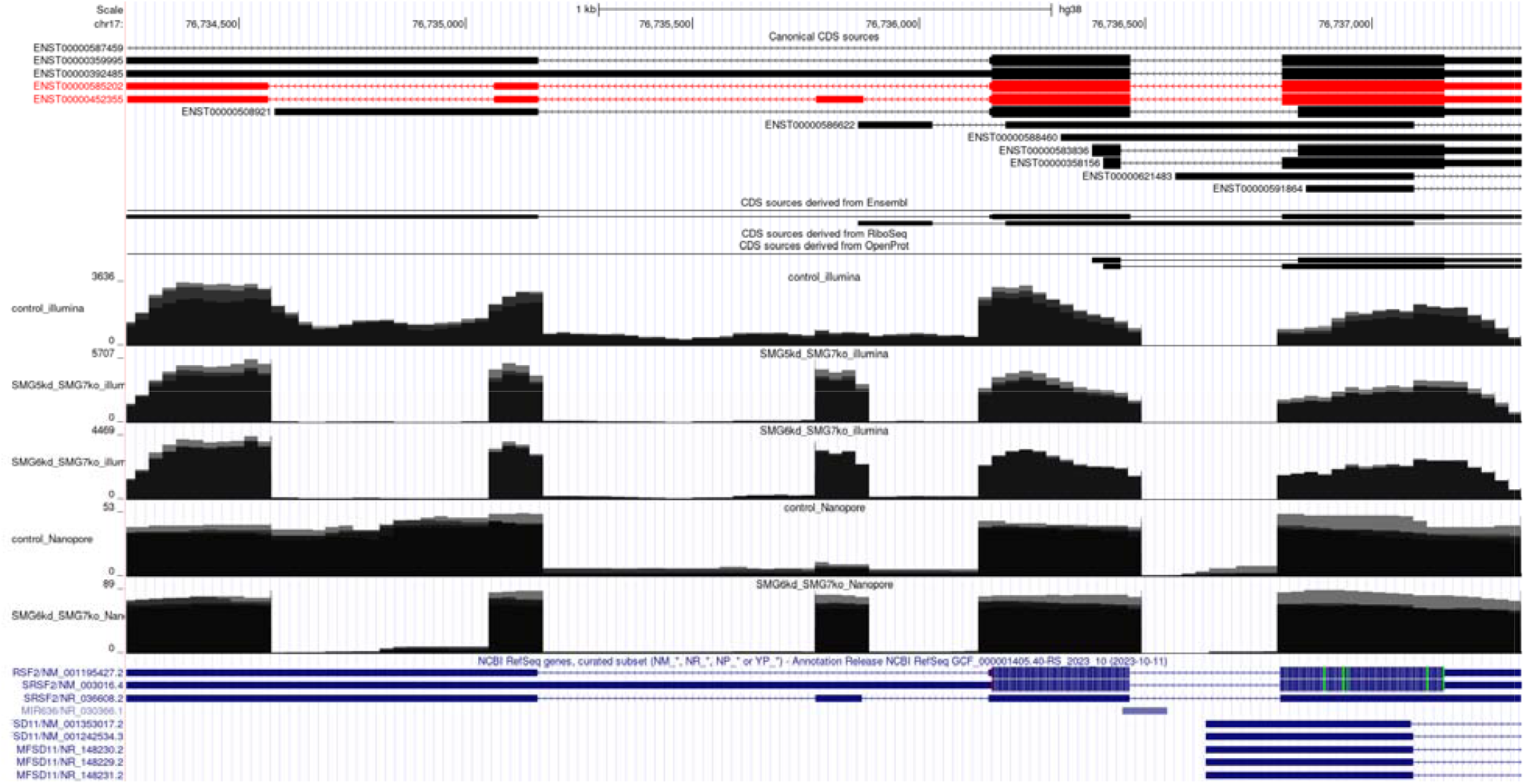
TrackHub view for the SRSF2 gene. The hub comprises the transcriptome assembly and coverage of selected RNA libraries that were aggregated. Transcript-to-CDS matches from the same source were grouped t facilitate interpretation. As part of the UCSF Genome Browser, this hub can be used in addition to any other hub, facilitating data integration.

Figure 7 showcases the onboarding process for NMDtxDB, initiated through a dedicated button on the sidebar’s lower section. The process is designed to teach users how to use each application components and how to interpret data analysis. It guides users to effectively explain the NMDtxDB’s features. The interface highlights interactive features like download options and selection input to enhance user navigation. Moreover, detailed explanations appear when hovering over table column headers. Finally, users can download tables and plots based on the current selected view.

**Figure 7.**
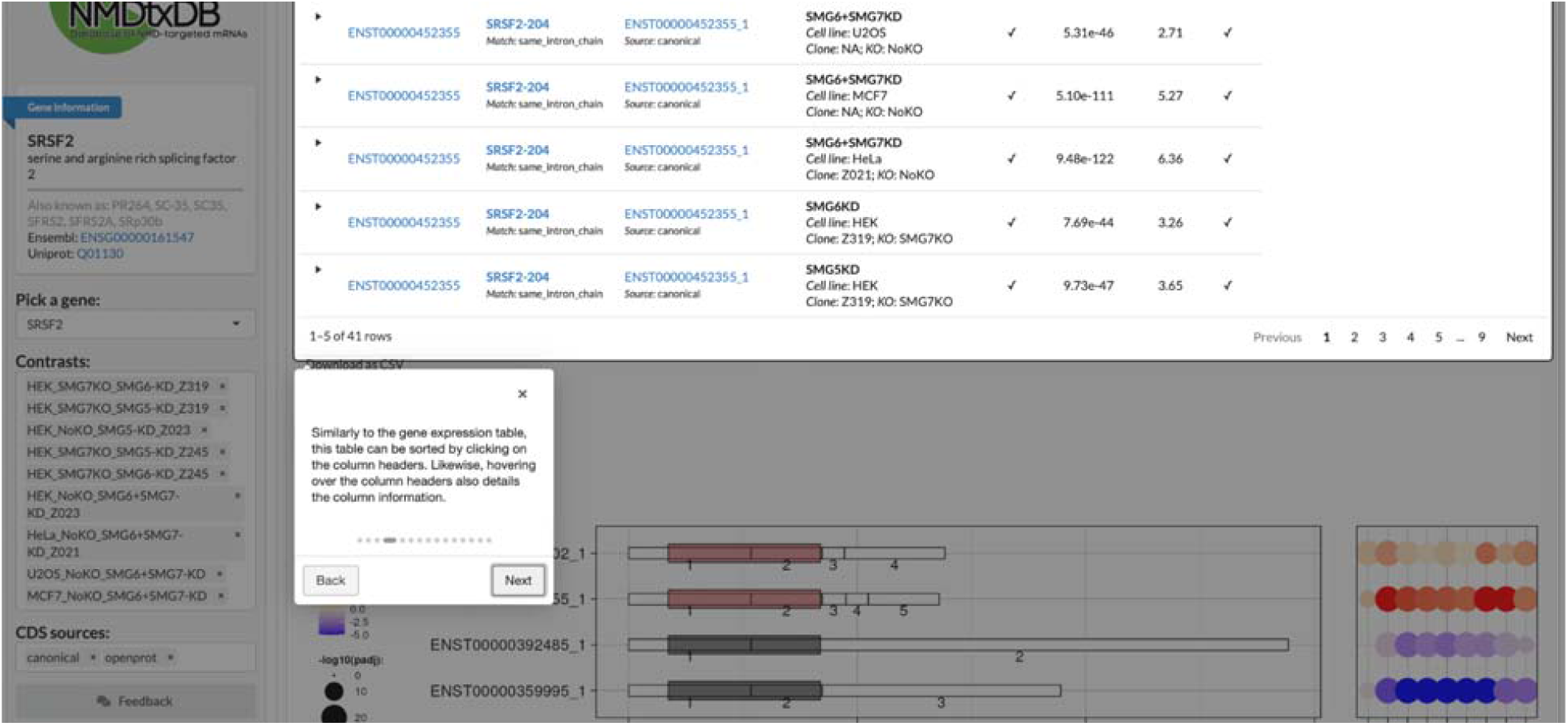
NMDtxDB tutorial. The tutorial serves as onboarding for new users. Each element of the user interface is highlighted and explained.

**Figure 8.**
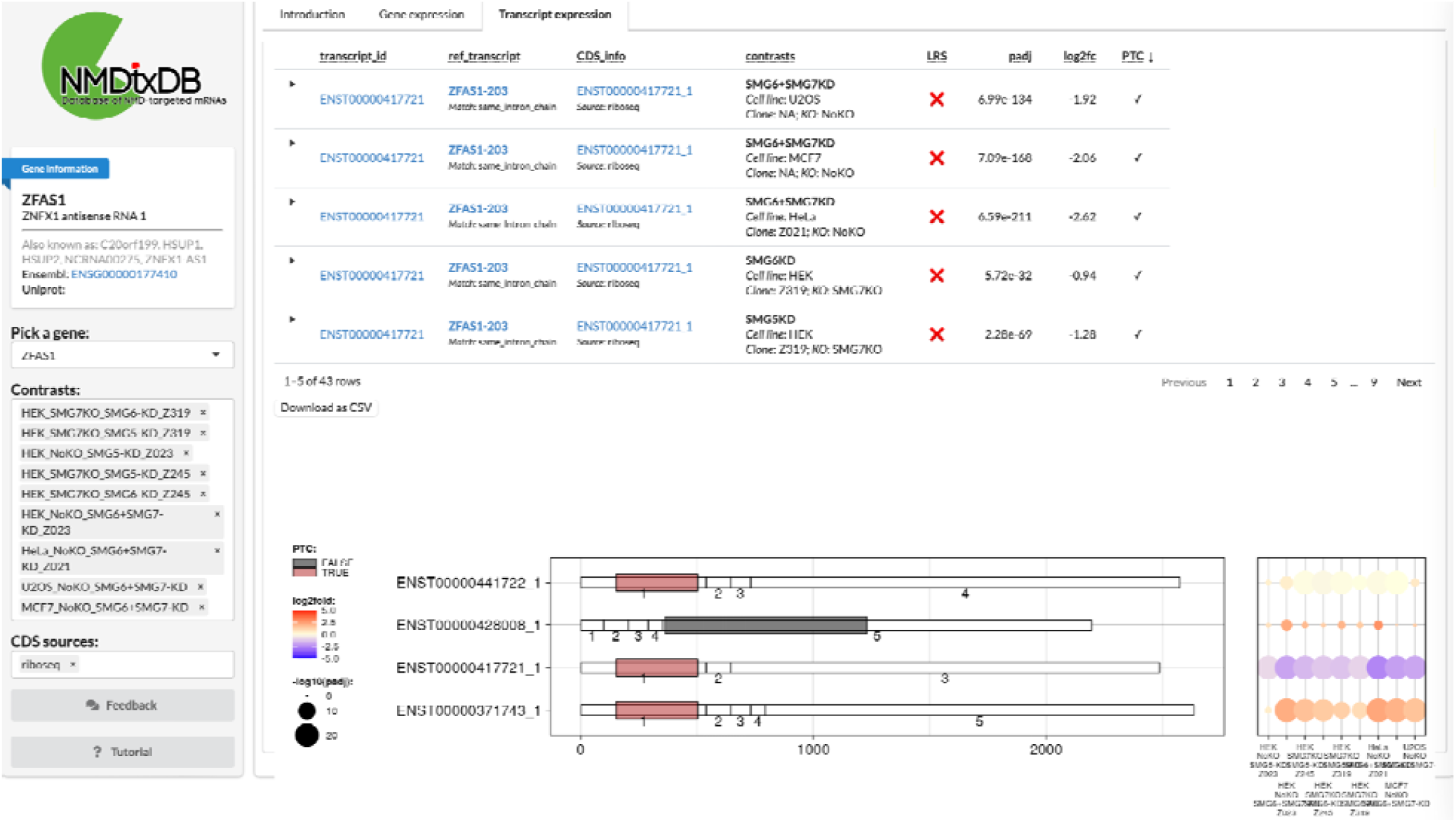
NMDtxDB page for ZFAS1, a known NMD target that is annotated as a non-coding gene. Th transcript ENST00000371743 is annotated as lncRNA, but there is evidence for translation from the consensus Ribo-seq resource. In addition, this transcript is up-regulated in multiple contrasts, as shown in the dot plot.

## Discussion

In this manuscript, we presented a dataset, a workflow, and a database providing an easy-to-access perspective on the effects of alternative splicing in context with the NMD-depletion. We used Illumina and Nanopore DRS to assemble a transcriptome of NMD factor depleted cell lines. In this work, we combine the two sequencing strategies to better characterize the NMD-sensitive transcriptome. For example, in NMDtxDB, we flag which transcripts have evidence from long-reads, providing information to the user to decide on which transcript to select for further analysis. A small-scale and PCR-based approach to sequence mRNA with long reads from cells depleted of NMD factors has been reported previously (25).

Second, we annotated CDS regions in our transcriptome based on two databases of coding sequences in addition to state-of-the-art Ensembl database: (1) OpenProt uses the PRIDE archive to find evidence for alternative proteins by re-analyzing published mass spectrometry datasets (16). (2) GENCODE Ribo-seq ORFs is a community-driven initiative that created a consensus set of open translation reading frames from seven Ribo-seq experiments (17). This expansion on coverage led to the discovery of 9,823 novel transcripts containing PTCs. Once the stop codons are experimentally validating, these transcripts can provide valuable insights into the NMD regulatory pathway by revealing potential novel NMD-sensitive transcripts that originate from alternative splicing events or transcripts annotated as noncoding.

A notable example for the utility of NMDtxDB is *ZFAS1*, a known NMD target that is currently annotated as a non-coding gene (6). Figure 9 shows the NMDtxDB page for *ZFAS1* and reports the up regulation for a transcript that has evidence of PTC based on the GENCODE Ribo-seq CDS.

**Figure 9.**
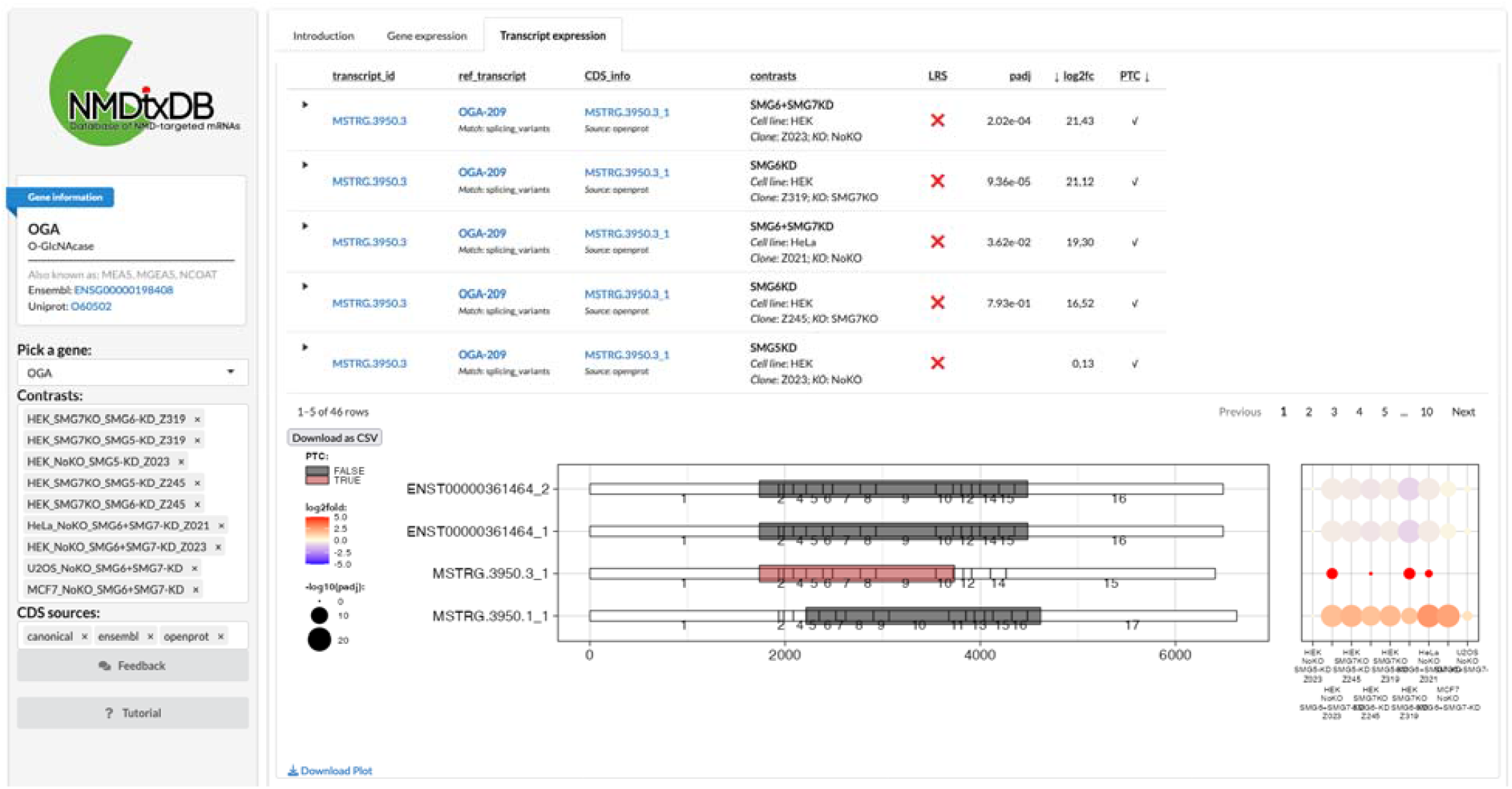
NMDtxDB page for OGA, a relevant metabolic enzyme that was recently described as an NMD target. The novel transcript MSTRG.3950.3 harbors a PTC derived from an OpenProt CDS source. This transcript is also significantly up-regulated in 3 conditions.

NMD depletion results in higher expression levels of transcripts harboring PTCs and, consequently, the genes comprising these transcripts. NMD activity is generally considered consistent across cells and tissues, yet its targets often include post-transcriptional regulators, which induce secondary effects in transcript expression. The expression changes in these regulators can alter the expression of downstream genes that do not contain PTCs. Combining PTC annotations with expression data from NMD-depleted cells allows for improved annotation of NMD-sensitive transcripts. It is essential to describe these targets to have a more complete overview of the NMD transcriptome, with potential future applications. For another example, Figure 9 illustrates the NMDtxDB page for *OGA*. The *OGA* enzyme plays a significant role in energy metabolism pathways and has recently been identified as an NMD target associated with human disease (26).

In summary, the NMDtxDB user interface is broadly accessible, simple to use and combines detailed information on novel, NMD-sensitive transcripts along with their PTC status. In the future, we plan to extend this unique web resource with new modalities of NMD evidence once they become available such as data from ribosome footprinting experiments or RNA fragments originating from endocleavage events (20,27,28). We also anticipate the use of machine learning methods to facilitate sorting NMD sensible from transcripts that undergo changes due to off-target effects, which could further improve user-decision making and biological interpretation.

Given the open-source nature of the database, NMDtxDB can serve as a platform for a further examination and access to NMD transcriptomes, with the common objective of dissecting the pathway and examining its impact on human health.

## Methods

### Cells lines, siRNA-mediated knockdown and knockout cells using CRISPR-Cas9

Cell line handling and siRNA-mediated knockdown preparation are consistent with our previous work (6). Flp-In-T-REx-293 (human, female, embryonic kidney, epithelial; Thermo Fisher Scientific), HeLa Tet-Off (human, female, cervical adenocarcinoma; Clontech), MCF7 (human, female, breast carcinoma; kind gift from Kay Hofmann, University of Cologne) and U2OS (human, female, osteosarcoma; kind gift from Kay Hofmann, University of Cologne) cells were cultured in high-glucose, GlutaMAX DMEM (Gibco) supplemented with 9% fetal bovine serum (Gibco) and 1x Penicillin Streptomycin (Gibco). The cells were cultivated at 37°C and 5% CO2 in a humidified incubator. The Flp-In-T-REx-293 SMG7 knockout cell lines were established and described previously (6).

Briefly, the CRISPR-Cas9 knockouts employed the Alt-R system (Integrated DNA Technologies) and reverse transfection of a Cas9:guideRNA ribonucleoprotein complex using Lipofectamine RNAiMAX. The construct targeting SMG7 target was previously described (6).

For siRNA-mediated knockdown, the cells were seeded in 6-well plates at a density of 3 × 10^5^ cells per well and reverse transfected using 2.5 μl Lipofectamine RNAiMAX and 60 pmol of the respective siRNA(s) according to the manufacturer’s instructions. In preparation for Nanopore direct RNA sequencing, 2.5 × 10^6^ cells were reverse transfected in 10 cm dishes using 6.25 μl Lipofectamine RNAiMAX and 150 pmol siRNA. The siRNA sequences were as follows: Luciferase 5′ □ CGUACGCGGAAUACUUCGA-3′; SMG5 5′ □ GAAGGAAAUUGGUUGAUAC-3′; SMG6 5′ □ GGGUCACAGUGCUGAAGUA-3′; SMG7 5′ □ CGAUUUGGAAUACGCUUUA-3′.

### RNA extraction, library preparation and sequencing

Cells were harvested 72h after siRNA transfection using peqGOLD TriFast (VWR Peqlab) and RNA was isolated following the manufacturer’s instructions, using 150 μl 1-Bromo-3-chloropropane (Molecular Research Center, Inc.) instead of 200 μl chloroform per 1 ml of TriFast. For Nanopore direct RNA sequencing, poly(A) enrichment was performed in two consecutive rounds using 100 μg of total RNA and 200 μl Dynabeads Oligo(dT)_25_ per sample (Thermo Fisher Scientific), following the manufacturer’s instructions.

### Illumina RNA Sequencing

The library preparation was performed with the TruSeq mRNA Stranded kit (Illumina). After poly-A selection (using poly-T oligo-attached magnetic beads), mRNA was purified and fragmented using divalent cations under elevated temperature. The RNA fragments underwent reverse transcription using random primers. This is followed by second strand cDNA synthesis with DNA Polymerase I and RNase H. After end repair and A-tailing, indexing adapters were ligated. The products were then purified and amplified to create the final cDNA libraries. After library validation and quantification (Agilent tape station), equimolar amounts of library were pooled. The pool was quantified by using the Peqlab KAPA Library Quantification Kit and the Applied Biosystems 7900HT Sequence Detection System and sequenced on an Illumina NovaSeq6000 sequencing instrument and a PE100 protocol.

### Nanopore Direct RNA Sequencing

Library preparation for Nanopore Sequencing was carried out using the Direct RNA Sequencing Kit (SQK-RNA002) from Oxford Nanopore Technologies following manufacturer’s guidelines. Sequencing was carried out on the GridION X5 platform as per manufacturer’s guidelines using R9.4.1 flow cells.

### RNA-sequencing read processing

Illumina reads pre-processing and alignment have been detailed before (6). Nanopore DRS reads were base called with guppy and aligned with minimap2:

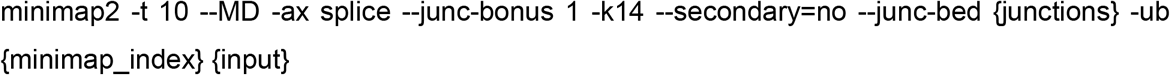

All RNA-seq datasets were aligned against the Ensembl genome GRCh38.p13 (v102). We only consider autosomes or sex chromosomes for alignment. All features in other chromosomal regions were discarded after alignment.

### Guided transcriptome assembly workflow

The workflow for identifying and annotating human NMD target transcripts relied on the snakemake workflow management system. Snakemake provides a framework for modular, scalable, reproducible workflows (29). The nmd-wf is available at https://github.com/dieterich-lab/nmd-wf under the MIT license. The transcriptome assembly was carried out in two stages, as only a subset of RNA samples were sequenced ONT-seq as well. In the first step, the reference transcriptome (Ensembl GRCh18 v102) was used as a guide to generate novel transcriptome annotations for pairs of samples with Illumina and Nanopore sequencing. StringTie was used with the parameters --mix and --rf to accomplish this. The merged GTF file from stage 1 was used as a guide to the second StringTie call for each Illumina sample. The resulting annotations were merged a last time. Annotation merge was conducted with stringtie -- merge and min_iso_prop=0.1 and min_cov=3 parameters. The first stage used the reference transcriptome, while the second stage used the first stage merge as the guide parameter (-G). The reference gene and transcript names were as well the class codes were obtained by running GffCompare against the reference annotation (30).

### Benchmarking assembly parameters versus CHESS

We optimized StringTie parameters by performing a grid search for a single sample (33G-30) and compare the assembly with GffCompare (30). Based on the F1 metric, the mix preset obtained the highest score for the exon and intron feature level, while the conservative metric obtained the top for the transcript feature. We note that in terms of F1 metric, the highest source of decrease in score was the lack of guide (noguide preset).

### Matching long reads to transcripts

We mapped the DRS reads to the transcriptome to match the reads to the assembled transcriptome. The read alignment was computed with:

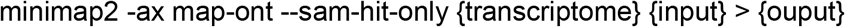

Following the read alignment, our approach involves selecting primary alignments for the computation of transcript coverage. Primary alignments with 90% sequence coverage on read level were considered as evidence of long read support for that transcript as defined by BamSlam (31).

### Coding sequence curation

The database has two classes of CDS matches: canonical or from sources. For annotated transcripts that have CDS matches to the EnsEMBL database, we simply imported this information to the database. In NMDtxDB this CDS source was named canonical.

To find matches for novel transcripts or transcripts without an annotated CDS, we apply a *de novo* discovery approach. This approach uses three databases of CDS: Ensembl, or Ribo-Seq ORFs and OpenProt, named ensembl, riboseq and openprot. First, we mapped the start codon from CDS sources from the genomic coordinates to the transcript coordinates. Next, we trimmed the transcript sequence from the start codon. We applied a modified version of the longorf.pl script (https://github.com/bioperl/bioperl-live/blob/master/examples/longorf.pl) to find the longest and complete, i.e., with a canonical start and stop codons, CDS of each transcript.

The coordinates of the two branches are then integrated. Duplicated CDS are dropped, and just one record was retained using the following ranking list: Ensembl, riboseq or openprot. The diagram in Figure 10 details this workflow.

**Figure 10.**
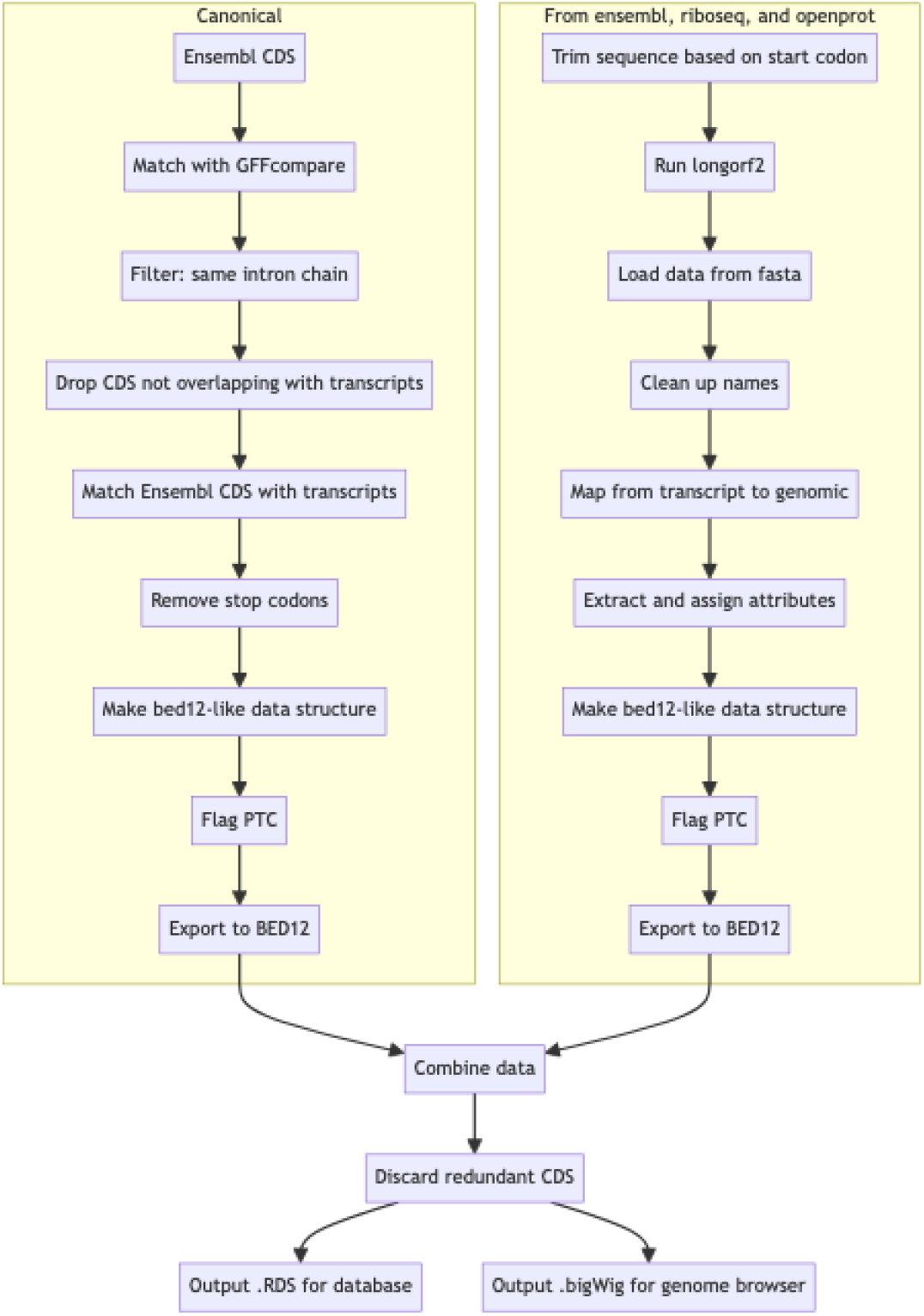
CDS integration workflow. The scheme shows the two branches of data processing for CDS integration. Database source are described in Table 2.

If multiple distinct CDS occur per transcript, they are reported. We compare these examples in Figure 11. It shows the canonical source as the primary source for CDS, followed by openprot. The combinations canonical+openprot, canonical+riboseq, canonical+ensembl, riboseq and ensembl provide a similar order of magnitude of CDS-to-transcript matches. Of note, the plot highlights a higher LR support for combinations with higher degree that comprise the canonical source, specifically columns 1 to 4. Upset plot was created with ComplexHeatmap packages (32).

**Figure 11.**
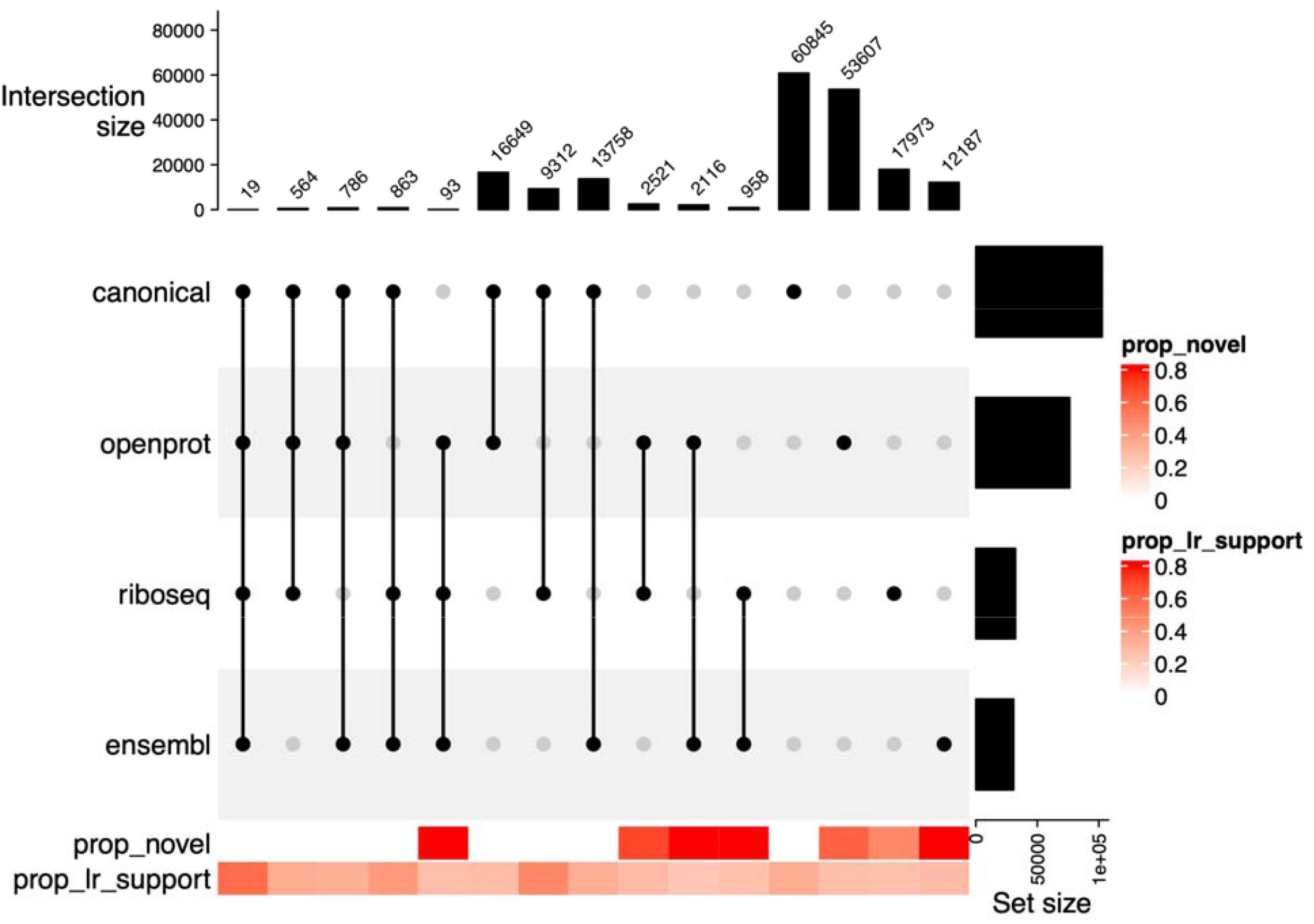
Distinct combination of CDS source per transcript. The plot compares the source of CDS per transcript, using combinations to show overlapping CDS sources per transcript. The color bar in the bottom shows, respectively, the proportion of novel transcripts or transcripts with LR support by combination.

### Gene and transcript expression modeling

Salmon was used to compute gene and transcript expression, which was then summarized with tximport / in the dtuScaledTPM mode (33,34). Transcripts were filtered using the DRIMSeq::dmFilter function with parameters of minimum gene expression of 10, minimum transcript expression of 1, and minimum transcript proportion of 0.10, resulting in 82,799 transcripts (16,812 genes). The remaining transcripts were discarded from further modeling. Gene counts were imported using tximport in the lengthScaledTPM mode; only genes containing transcripts that passed the transcript filtering were considered for downstream analysis.

The DEXSeq package was used to perform differential transcript usage analysis (35). The experiment was run with a treatment versus control design, as per the contrast listed in Table 1. The full model is defined as follows: ∼ sample + exon + group:exon. The reduced model, which is contrasted against the full model, is defined as ∼ sample + exon.

The DEseq2 package with Independent Hypothesis Weighting (IHW) correction was used to analyze the differentially expressed genes (36,37). Genes were filtered using a minimum count threshold, with the row sum of counts in the dds object having to be greater than or equal to 10 in at least three samples. The lfcShrink function from the ashr (Adaptive Shrinkage) package was used to obtain shrunken fold changes (38).

### Database and web application

The web-application infrastructure is based on our previous work (39). In short, the Shiny web-application is hosted via Open Analytics ShinyProxy server, in the demilitarized zone of the high-performance computer cluster at Heidelberg University Hospital’s Klaus Tschira Institute for Computational Cardiology. The application is built with the Golem framework (https://github.com/ThinkR-open/golem), using ggplot2 (https://ggplot2.tidyverse.org), and ggtranscript (40) for visualization. In contrast to our previous application, we replaced the PostgreSQL database by a docker checkpoint-restore system that improves user experience by reducing application loading times. Gene card information is obtained via the mygene REST API (41).

## Data availability

Raw files for the RNA-sequencing are available at the Sequence Read Archive at https://www.ncbi.nlm.nih.gov/sra/PRJNA1054031. An archive of source code and database are available at and https://zenodo.org/records/10533763.

## Code availability

Code for workflow and web-application are available at https://github.com/dieterich-lab/nmd-wf and https://github.com/dieterich-lab/NMDtxDB (MIT license).

## Acknowledgements

We would also like to thank Harald Wilhelmi for support with infrastructure and user experience optimization. Federico Marini for feedback on the user interface. Member of the Dieterich lab and anonymous testers for suggestions. We also thank the Cologne Center for Genomics, CCG, for preparing the sequencing libraries and operating the Illumina sequencer. Nanopore Next-Generation Sequencing was carried out at the West German Genome Center Düsseldorf.

